# Rapid diversification of a natural Heterosigma akashiwo virus population during a host bloom

**DOI:** 10.64898/2026.02.06.704369

**Authors:** Jun Xia, Lingjie Meng, Yue Fang, Hiroki Ban, Yusuke Okazaki, Takashi Yoshida, Hisashi Endo, Keizo Nagasaki, Hiroyuki Ogata

## Abstract

Despite the ecological importance of viruses, our understanding of their evolutionary dynamics in natural environments remains limited. This gap is particularly pronounced for giant dsDNA viruses of the phyla *Nucleocytoviricota* and *Mirusviricota*. Most knowledge on their population genetic dynamics is derived from a small number of laboratory-based experiments, whereas patterns in nature are rarely observed. To overcome this limitation, we traced genetic structure and transcription status of Heterosigma akashiwo virus (HaV) using high-frequency, time-resolved sampling during a host bloom in a coastal area of Japan, by integrating cell counting, metabarcoding, metagenomic and metatranscriptomic sequencing. Our study revealed that HaV dominated the giant virus community in most samples, with the relative abundances of up to 56%. Despite the high abundances, the HaV population exhibited a relatively low level of microdiversity but with a high pN/pS ratio compared to other giant viruses in the study site. Microdiversity increased during the early sampling period, reached a maximum at mid-sampling, and decreased during the later period, consistent with rapid diversification during viral expansion, possibly driven by both *in situ* mutations and the succession of pre-existing minor variants. Several accessory genes, including a glycosyltransferase and an endonuclease, were highly expressed, providing functional evidence consistent with host interaction-driven selective pressure during the bloom. Together, these results indicate that HaV population dynamics during algal blooms are shaped by host-driven selection acting on standing genetic variation.

## Introduction

Viruses usually evolve much faster than cellular organisms (Duffy *et al*., 2008; Holmes, 2010). Among viruses, giant viruses (GVs) generally exhibit lower mutation rates than smaller viruses (e.g., ssDNA and RNA viruses), albeit being still higher than those of cellular organisms (Sanjuán *et al*., 2010; Duchêne and Holmes, 2018). GVs are large double-stranded DNA (dsDNA) viruses, typically in *Nucleocytoviricota* (Aylward *et al*., 2021) and *Mirusviricota* (Gaïa *et al*., 2023), that infect a wide range of eukaryotes (Iyer *et al*., 2006). Their large genomes are partly due to frequent horizontal gene transfer (HGT) from cellular organisms and viruses (Irwin *et al*., 2022; Wu *et al*., 2024; Yi *et al*., 2024), which enhances adaptive evolution through the acquisition of auxiliary metabolic genes modulating host cellular metabolism during infection (Ha *et al*., 2021; Brahim Belhaouari *et al*., 2022; Meng *et al*., 2023b). Intra-species diversity (i.e., microdiversity) may also be important for the adaptation of GVs. A recent study revealed that in the Uranouchi Inlet, marine GVs with persistent presence across seasons tend to exhibit higher levels of microdiversity than those showing sporadic appearance, suggesting that larger effective population size enhances the fitness of GVs under the virus-host arms race during prolonged interactions with their hosts (Fang *et al*., 2025). However, short-term and high-resolution evolutionary dynamics of GVs in natural environments remain largely uncharacterized.

Tracking short-term evolutionary dynamics of natural GV communities is challenging, because the community changes more through advection and diffusion of water masses than through viral evolution. The Uranouchi Inlet is an enclosed coastal region located in Kochi Prefecture, Japan. Harmful algal blooms (HABs), which can be easily tracked, occur frequently in the Uranouchi Inlet (Takahashi *et al*., 2021). The bloom events are accompanied by dramatic changes not only in plankton communities but also in viral communities (Vincent *et al*., 2023). Thus, this inlet provides an ideal environment for observing the evolutionary dynamics of plankton and viral communities with minimal effects of water mass changes (Prodinger *et al*., 2021; Fang *et al*., 2025). One of the major HAB species in the Uranouchi Inlet is the raphidoflagellate *Heterosigma akashiwo*, belonging to family Chattonellaceae and often blooming in late spring (Prodinger *et al*., 2021; Funaoka *et al*., 2023).

Some GVs are known algal bloom regulators that can shape microbial community structures and directly or indirectly influence the duration and intensity of blooms by killing blooming algae (Nagasaki *et al*., 1994b; Brussaard *et al*., 1996; Jacquet *et al*., 2002; Gajigan *et al*., 2025). Heterosigma akashiwo virus (HaV; family *Phycodnaviridae*, genus *Raphidovirus*, species *Raphidovirus japonicum*) is a GV that infects *H. akashiwo* and is known to have a role in terminating the HAB through its lytic infection (Nagasaki *et al*., 1994a; Nagasaki *et al*., 2002). The sporadic boom-and-bust population dynamics of *H. akashiwo* and associated HaV make this alga-virus system an intriguing model for understanding their co-evolutionary processes. A recent cross infection study involving 60 *H. akashiwo* stains and 22 HaV strains revealed highly diverse infection spectra, ranging from strain-specific to broad host-range types (Funaoka *et al*., 2023), suggesting the existence of genetic diversity in both host and virus populations that determine the infection specificity. Such rapid shifts in viral genetic structure can emerge not only from the accumulation of *in situ* mutations but also from the dynamic succession of pre-existing variants; specifically, minor variants that persist at low frequencies may rapidly expand and become dominant when their specific host strains proliferate (Roux *et al*., 2017; Zhou *et al*., 2025). Therefore, accounting for the dynamics of this standing microdiversity is crucial for a comprehensive understanding of viral adaptive processes during bloom events. However, like their host, HaVs show a sporadic pattern of proliferation dynamics, which may lead to a reduced level of genetic diversity due to recurrent population bottlenecks. Currently, there is no study that investigated the level of genetic diversity and dynamics of a natural HaV population.

To investigate the diversity of HaV at the population level, we conducted high-frequency, daily sampling at the Uranouchi Inlet during a *Heterosigma akashiwo* bloom period. We successfully recovered a near complete environmental HaV genome together with other GV genomes through metagenomics. By integrating these time-resolved metagenomic data with cell count, 18S rDNA metabarcoding, and metatranscriptomic data, we characterize the fine-scale temporal dynamics the genetic structure and gene expression of a HaV population in the inlet, which provide insights into the eco-evolutionary dynamics of HaV and other GVs during algal blooms.

## Materials and Methods

### Sampling

Seawater samples were collected at two sites in Uranouchi Inlet: A-site in red tide area of innermost bay (133.40°E, 33.43°N) and B-site near a fishery farm located in the middle of the inlet (133.36°E, 33.41°N) (Fig. S1). Sampling was conducted from May 30 to June 7, 2019—three consecutive days at A-site (June 3 to 5 or Day-5 to Day-7) and nine consecutive days at B-site (May 30 to June 7; Day-1 to Day-9) (Table S1). Samplings were performed during the daytime (08:20–13:50). Samples of A-site were collected from clearly observable *Heterosigma akashiwo* blooming patches.

At A-site, surface and subsurface chlorophyll maximum (SCM; 2.5–3.1 m) layers were sampled, while at B-site SCM (1.0–3.1 m) samples were collected. Seawater was prefiltered through a 144 µm mesh to remove large particles. For each metagenomic (metaG) sample, 10 L of seawater was filtered sequentially through 3 µm membrane filters and four 0.22 µm Sterivex units (three 3–144 µm and fourteen 0.22–3 µm samples were used in downstream analysis). For each metatranscriptomic (metaT) sample, 500–1,000 mL seawater was filtered through 3 µm polycarbonate filters, and filters were preserved in RNAlater and stored at -80°C (nine 3–144 µm samples were used in downstream analysis). For each 18S rDNA metabarcoding (metaB) sample, 500–1,000 mL of seawater was filtered through 3 µm filters (fourteen 3–144 µm samples were used in downstream analysis). For each sample for microscopic cell count, 2 L of unfiltered surface or SCM seawater was fixed with Lugol’s solution (2%, v/v) and stored at 4°C.

### DNA and RNA extraction

DNA and RNA were extracted following the previously described protocols (Endo *et al*., 2018). Purified DNA was dissolved in 30 µL of low-TE buffer for metabarcoding (metaB) analysis and in 30 µL of Milli-Q ultrapure water for metaG analysis. For RNA extraction, samples were mixed with β-mercaptoethanol, RLT lysis buffer, and glass beads, followed by bead beating and centrifugation. The supernatant was processed with the RNeasy Mini Kit (QIAGEN), and residual DNA was removed using RNase-Free DNase. Extracted DNA and RNA were stored at −20°C until further use. DNA and RNA concentration and purity (A260/280) were quantified using a Qubit fluorometer (Thermo Fisher Scientific), a Nanodrop spectrophotometer (Thermo Fisher Scientific) and a TapeStation system (Agilent Technologies).

### Library construction, sequencing and data processing

MetaG libraries were prepared using TruSeq Nano DNA Kit and sequenced on Illumina Miseq 300PE platform.

MetaT libraries were prepared using the NEBNext® Poly(A) mRNA Magnetic Isolation Module, NEBNext® UltraTMII Directional RNA Library Prep Kit, and NovaSeq 6000 150PE sequencing was conducted.

For metaB analysis, the V8–V9 regions of the eukaryotic 18S rDNA gene was amplified using primers V8f–1510r (positions 1,422–1,797 in *Saccharomyces cerevisiae* gene) following the previous method (Bradly *et al*., 2016; Prodinger *et al*., 2020). PCR products were checked by agarose gel electrophoresis, purified, and diluted with 25 µL of ultrapure water. MetaB libraries were constructed using MiSeq Reagent Kit v2 and sequenced on Illumina Miseq 300PE platform. MetaB reads were processed with QIIME2 (v2020.2) (Bolyen *et al*., 2019). Trimming, quality filtering, denoising, merging, dereplication, chimera removal, and generation of amplicon sequence variants (ASVs) were performed using DADA2 plugin integrated in QIIME2 (q2-dada2 version 2020.2.0) (Callahan *et al*., 2016). Taxonomic classification was performed using the PR2 database (version 4.14.0) (Guillou *et al*., 2012) with BLASTn (≥97% identity). After excluding ASVs classified as metazoa or fungi as well as singleton ASVs, sequence data were subsampled to the minimum size among all 14 samples to compute relative abundances of ASVs.

After quality filtering, sequencing yielded 2.09 million metaB paired reads (1.26 Gbp, N=14), 2.56 billion metaG reads (725.75 Gbp, N=17), and 0.65 billion metaT reads (195.21 Gbp, N=9) (Table S4).

### Reconstruction of GV-MAGs

The pipeline hedera (version 0.0.5) (https://github.com/banhbio/hedera) developed in previous studies (Fang *et al*., 2025; Liu *et al*., 2025) was improved for the generation of GV metagenome-assembled genomes (GV-MAGs). The complete workflow from raw reads processing to final GV-MAG construction is described and illustrated in Supplementary Text and Fig. S2. Briefly, after co-assembly of metaG reads from 17 samples (three 3–144 samples and fourteen 0.22–3 samples), 2,515,850 contigs (≥1 kbp, mean length 2,796 bp) were generated, of which 590,170 contigs (≥2.5 kbp, mean length 6,910 bp) were retained for binning. Raw bins (1,794 bins) produced by MetaBAT2 using differential coverage across all 17 metagenomic samples, as well as 1,734 unbinned contigs (≥30 kbp) were included for the downstream analysis. Based on the GV marker gene density index (>5.75) (Fang *et al*., 2025), 432 bins and 65 single contigs were retained. None was removed during the GV contig assessment. After delineation and second decontamination using Anvi’o (Eren *et al*., 2015), 504 GV-MAG candidates were retained. One contig was later removed from a bin due to the presence of abundant prokaryotic core genes. Subsequent manual delineation, phylogeny-informed MAG assessment (PIMA) and dereplication (no MAG was duplicated at 95% ANI threshold) yielded 424 high-quality GV-MAGs (59.3 kbp–1.36 Mbp), including a MAG corresponding to a HaV population.

Additional manual curation and refinement of the HaV-MAG were conducted. All contigs obtained from metagenomic co-assembly were aligned to the reference genome of the isolate HaV53 (Accession: NC_038553.1) (Ogura *et al*., 2016) using BLASTn (best hit, E-value<1e-5). Contigs showing reliable alignments were identified as potential HaV-contigs. Reads mapping to these contigs were extracted from sorted BAM files with Samtools (Li *et al*., 2009) and then reassembled to final HaV-contigs using SPAdes (v3.15.4) (Nurk *et al*., 2017) with “--meta” option. The reassembled HaV-contigs were reordered to form the final HaV-MAG based on HaV53 reference genome by GenomeNet’s ReCCO (v1.0-beta) (https://www.genome.jp/ftp/tools/ReCCO/ReCCO_Readme.html). The original automatically generated HaV-MAG was replaced with this refined MAG for the following analyses.

### Sequence comparison, phylogeny and function analysis

Basic statistics of MAGs were obtained using SeqKit (v2.3.0) (Shen *et al*., 2016). Average nucleotide identity (ANI) between genomes was calculated with FastANI (v1.33) (Jain *et al*., 2018). Dereplication at 95% ANI was performed among the GV-MAGs generated in this study and the genomic data in the GOEV database (Gaïa *et al*., 2023) using dRep (v3.4.2) (dRep dereplicate options: -sa 0.95 -l 1000 -comp 50 --S_algorithm ANImf -nc 0.5 -N50W 0 -sizeW 1 --ignoreGenomeQuality) (Olm *et al*., 2017). Amino acid identity between HaV-MAG and HaV53 was assessed by BLASTp (E-value<1e-3).

A maximum-likelihood phylogeny of nucleocytoviruses (nucleocytoplasmic large DNA viruses, NCLDVs) was reconstructed using IQ-TREE2 (v2.2.2.6) (Minh *et al*., 2020) with the LG+I+F+G4 model based on a concatenated alignment of seven conserved markers (DEAD/SNF2-like helicase [SFII], DNA-directed RNA polymerase alpha subunit [RNAPL], DNA polymerase family B [PolB], Transcription initiation factor IIB [TFIIB], DNA topoisomerase II [TopoII], Packaging ATPase [A32], and Poxvirus Late Transcription Factor VLTF3) generated by ncldv_markersearch.py (v1.1) (Aylward *et al*., 2021). Mirusvirus major capsid protein HK97 sequences were identified with hmmsearch (E-value<1e-5) (v3.4) (Finn *et al*., 2011), aligned with MAFFT (v7.525) (Katoh and Standley, 2013), trimmed by trimAl (v1.5.0, “-gt 0.1”) (Capella-Gutiérrez *et al*., 2009), and used to construct a maximum-likelihood tree using IQ-TREE2 under the LG+R6 model. All trees were visualized in Anvi’o (v7.1) (Eren *et al*., 2015). Relative evolutionary divergence (RED) values were calculated to determine the family and order levels clades of GVs.

Gene-level synteny between HaV-MAG and HaV53 reference genome was determined by DiGAlign (similarity search used BLASTn, E-value<1e-2 were shown in green colors) (v2.0) (Nishimura *et al*., 2024). The HaV-MAG genomic structure was visualized using the R package gggenes based on GFF annotations. Multiple sequence alignment of the HaV major capsid protein was performed using Clustal Omega (v1.2.4) (Sievers *et al*., 2011). Protein 3D structures were predicted using AlphaFold3 (Abramson *et al*., 2024) via the AlphaFold Server and rendered in ChimeraX (v1.9) (Meng *et al*., 2023a).

GV ORFs predicted by prodigal were annotated by hmmsearch (E-value<1e-5) against GVOG (Aylward *et al*., 2021), 149 core NCVOGs (Yutin *et al*., 2009; Gaïa *et al*., 2023), EggNOG (euk+arc+bac+vir) (Hernandez-Plaza *et al*., 2023) and Pfam-A (Mistry *et al*., 2021). Annotation priority followed: (I) GVOG annotated by NCVOG; (II) GVOG annotated by EggNOG; (III) GVOG annotated by Pfam; (IV) 149 NCVOG annotation; (V) EggNOG annotation; (VI) Pfam-A annotation. If a higher-priority annotation was labeled as “unknown,” “unannotated,” “NA,” blank, or equivalent, the next level of annotation was adopted.

### Community and population analysis

Qualified metaG reads from each sample were mapped to 2,241 GV genomes (424 GV-MAGs from this study and 1,817 GOEV genomes) using minimap2 (Li, 2018) in CoverM (v0.7.0) (Aroney *et al*., 2025) with thresholds of >95% identity and >75% length of reads. The relative abundance of GV communities was estimated as TPM values and normalized to 100% per sample.

Diversity indices (Simpson, Shannon, and Pielou’s J) and Bray–Curtis dissimilarities were calculated using the vegan R package. ANOSIM and Mantel tests were performed with vegan and ade4, respectively. Non-metric multidimensional scaling (NMDS) was conducted using monoMDS (k=3), and the result was visualized with ggplot2.

For microdiversity analysis, simple repeats in GV-MAGs were masked (replaced with “N”) using RepeatMasker (v4.1.5) to avoid inclusion of ambiguous mapping of reads (Tarailo-Graovac and Chen, 2009). Qualified metaG reads from each sample were mapped to the repeat-masked GV-MAGs with bowtie2 (v2.5.0) (Langmead and Salzberg, 2012). Average depth for each gene was calculated based on sorted BAM files using samtools (Li *et al*., 2009). Microdiversity, including single nucleotide variants (SNVs), nucleotide diversity (ND), and pN/pS was assessed for each MAG in every sample using inStrain (v1.5.1) (Olm *et al*., 2021) with “-f 0.01” parameter, based on genes of repeat-masked GV-MAGs. ORFs with average depth≥10 and at least 25 SNVs were retained for analysis of pN/pS ratios.

Additional ND, which is denoted as Nei’s π in this study, and fixation index (F_ST_) for HaV-MAG were also calculated for the B-site samples (depth ≥10, repeats masked) using previously described methods (Sjöqvist *et al*., 2021; Fang *et al*., 2025) instead of inStrain. We used Nei’s π and F_ST_ for the comparison across samples, as these indices better control the compared nucleotide sites irrespective of the mapping depth differences across samples.

### Metatranscriptome analysis

MetaT reads (N=9, 3–144 µm size-fraction) were quality-filtered by fastp (v0.23.4) (Chen *et al*., 2018). Ribosomal RNA sequences were removed by searching qualified paired reads against the SILVA 138.1 database (Quast *et al*., 2013) (LSURef_NR99, SSURef_NR99, RF00001 and RF00002) using SortMeRNA (v4.3.6) (Kopylova *et al*., 2012). Transcripts per million (TPM) were calculated for each sample with CoverM (Aroney *et al*., 2025) against the 424 GV-MAGs derived from metaG data. TPM values were normalized to 100% per sample to represent relative expression abundance.

### Accession numbers

All reads generated in this study were deposited to the DNA Data Bank of Japan (DDBJ) under the bioproject PRJDB37393, biosample number from SAMD01619950 to SAMD01619977.

### Data availability

Giant virus MAGs generated in this study and related data files (424 GV-MAGs genome files, gene files, phylogenetic tree files, hmm profiles and summarized table for inStrain output) are accessed on GenomeNet at https://www.genome.jp/ftp/db/community/Uranouchi_GVMAGs_9days/.

## Results

### Abiotic and biotic conditions during the Heterosigma akashiwo bloom

Blooms of *Heterosigma akashiwo* are known to occur recurrently in spring in the Uranouchi Inlet. A red tide patch was visually observed at the surface of A-site on Day-5, indicating the bloom peak, while routine daily sampling was conducted at B-site and additional samples targeting the patchy bloom were collected around A-site. Environmental parameters were measured for all samples at B-site (from B_Day1_SCM to B_Day9_SCM), and part of A-site (A_Day5_S, A_Day7_S and A_Day7_SCM) (Table S1). Seawater temperature ranged from 22.28°C to 24.20°C. Chlorophyll *a* (Chl-*a*) concentrations fluctuated over nine days in the SCM layer at B-site (13.21-78.36 µg/mL). It exceeded the detection limit (99.99 µg/mL) at the surface of A-site on Day-5, and declined on Day-7 (2.91 µg/mL at surface, 15.36 µg/mL at SCM). Chl-*a* concentration was positively correlated with normalized DO, phosphate, and particulate organic carbon (POC) concentrations (Pearson’s correlation, adjusted *p* < 0.05; Table S2).

A total of 40 phytoplankton taxa were identified through microscopic cell count (Fig. 1A, Table S3). The two highest phytoplankton cell densities were observed in the samples A_Day5_S (15,816 cells/mL) and A_Day6_S (31,358 cells/mL). These samples were dominated by *H*. *akashiwo* (A_Day5_S, 8,464 cells/mL, 53.5%; A_Day6_S, 20,611 cells/mL, 65.7%). On Day-7, the cell density at A-site declined to 2,445 cells/mL in the surface, while it was comparable between Day-6 and Day-7 in the SCM layer (Day-6, 1,437 cells/mL; Day-7, 1,428 cells/mL). At B-site, phytoplankton abundance in the SCM layer exhibiting a fluctuating upward trend until reaching a peak at Day-7 (9,134 cells/mL) and decreased in the following two days. *H*. *akashiwo* cell density at B-site was low (average 58 cells/mL).

**Fig. 1.**
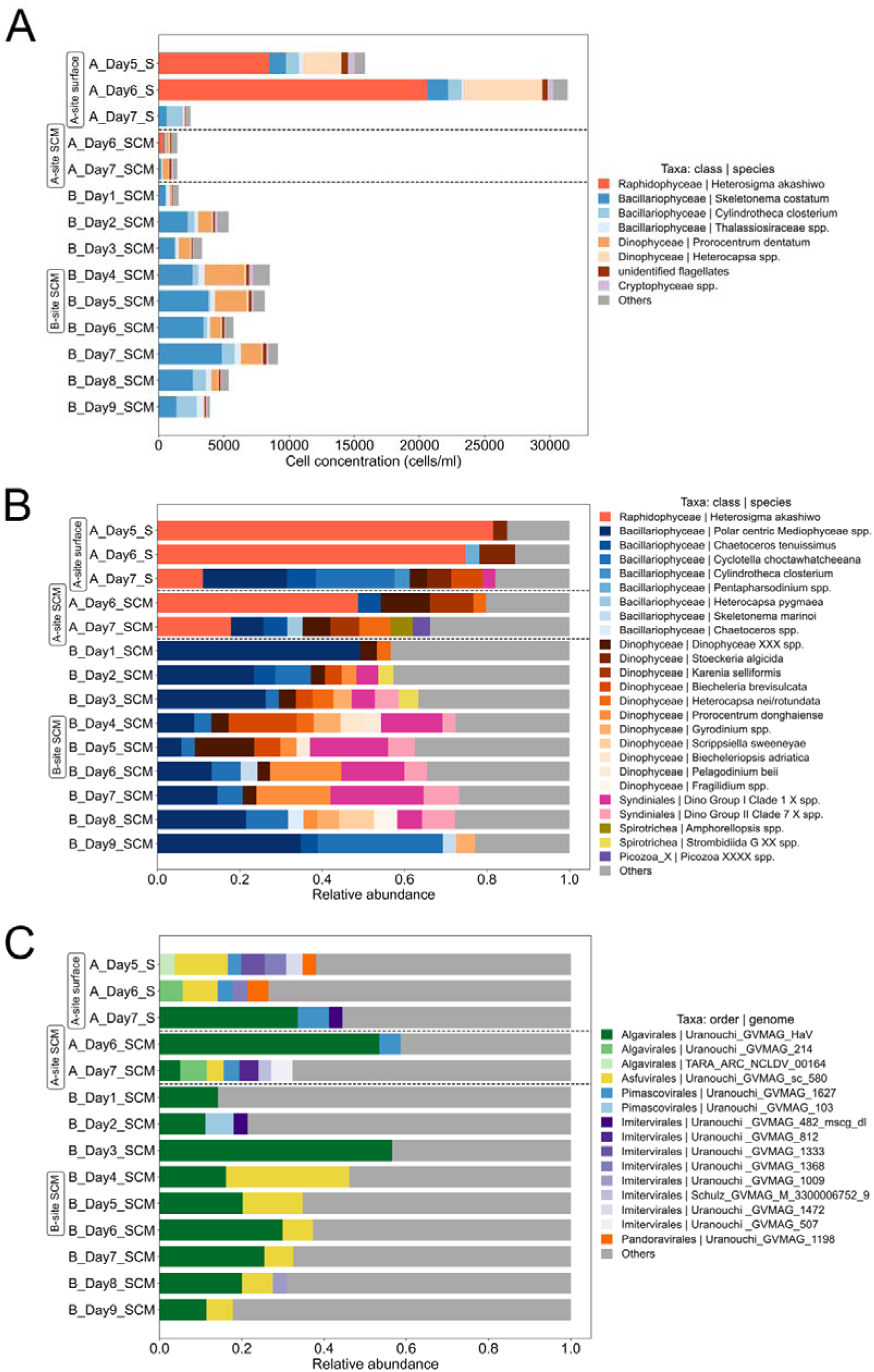
(A) cell counts of phytoplankton (taxa with >100 cells/mL in each sample average were shown); (B) relative abundance of microbial eukaryotic 18S community (metazoa and fungi removed, taxa relative abundance >3% for each sample were shown); (C) relative abundance of giant virus community (genome relative abundance >3% for each sample were shown).

A total of 1,050 non-singleton 18S rDNA ASVs (excluding metazoa and fungi) were recovered, of which 589 were taxonomically classified (Fig. 1B, Table S5). Consistent with the cell counting results, *H. akashiwo*, represented by seven ASVs, dominated the microeukaryote communities in the Day-5 and Day-6 samples at A-site (48.8–81.5%). Due to the dominance of *H*. *akashiwo,* these samples showed relatively low levels of 18S community diversity (Fig. S3A, Table S8). The relative abundance of *H*. *akashiwo* decreased on Day-7 at A-site (surface 11.0%, SCM 17.9%), while it was continuously low (<3%) in the B-site samples. Bacillariophyceae (diatoms), Dinophyceae (dinoflagellates), and Syndiniales (symbiotic dinoflagellates) were found to be abundant at B-site (Fig. 1B).

Eukaryotic community structures were significantly different between A-site (N=5) and B-site (N=9) (ANOSIM, *r*=0.934, *p*<0.001) (Fig. 2A). The community structures showed relatively large differences across A-site samples, except for A_Day5_S and A_Day6_S, which were collected from the bloom patch and displayed highly similar community compositions. At B-site, the community structures gradually changed along the sampling days from Day-1 to Day-9.

**Fig. 2.**
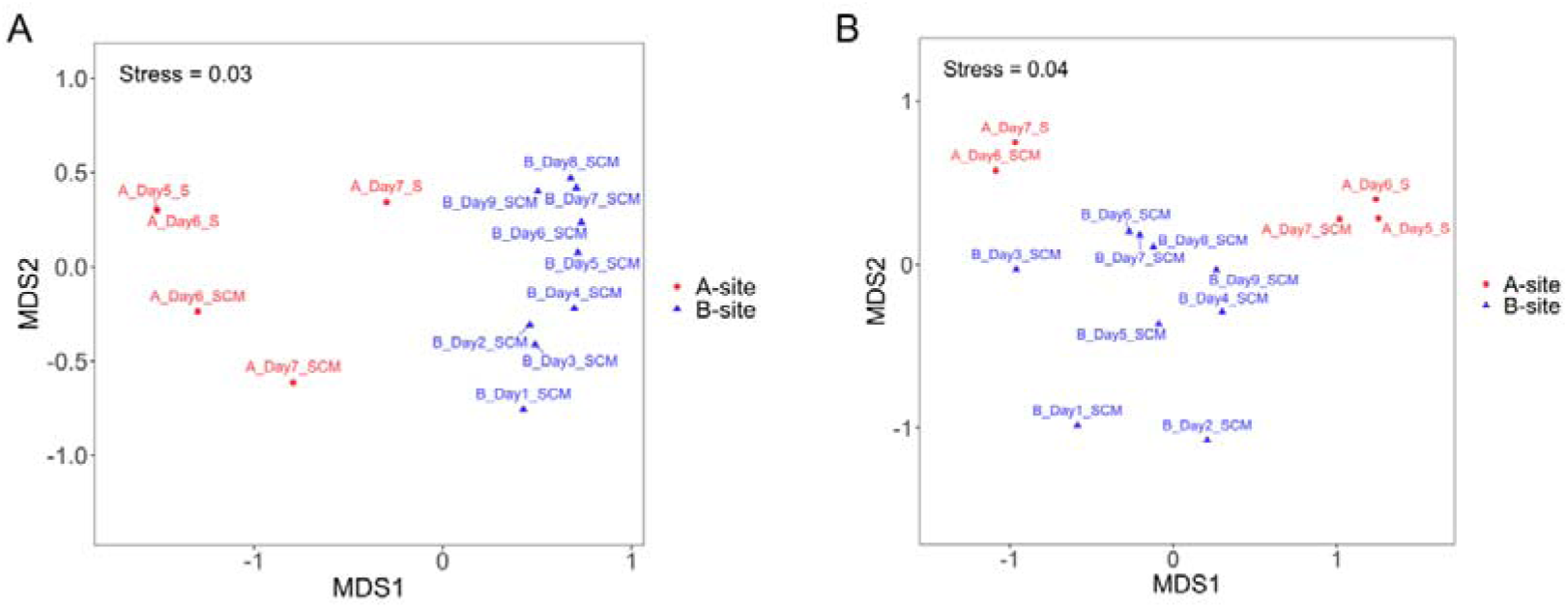
A nonmetric multidimensional scaling (NMDS) plot showing the diversity of eukaryotic and giant virus communities based on the Bray-Curtis dissimilarity. (A) microbial eukaryotic community (18S ASV level, 3–144 µm size-fraction). (B) giant virus community.

### Giant virus MAGs

Our computational pipeline and manual curation yielded 424 high-quality GV-MAGs with genome lengths ranging from 59.3 kbp up to 1.36 Mbp (see Materials and Methods). These MAGs were classified in either *Nucleocytoviricota* (420 GV-MAGs) or *Mirusviricota* (4 GV-MAGs). One MAG (HaV-MAG after manual curation) showed 99.86% ANI with the reference HaV53 genome. A phylogeny of nucleocytoviruses based on the core genes together with 1,692 genomes from GOEV assigned the 420 nucleocytovirus MAGs to *Imitervirales* (N=319), *Algavirales* (N=37), *Pandoravirales* (N=30), *Pimascovirales* (N=19), or *Asfuvirales* (N=14) (Fig. 3A). One nuclecytovirus MAG formed a long branch in the tree near the *Pandoravirales* clade and could not be assigned to a known order. A mirusvirus phylogenetic tree based on HK97 sequences together with those from 84 GOEV mirusvirus genomes assigned three mirusvirus MAGs to clade MR_01 and one to clade MR_02 (Fig. 3B). Except HaV-MAG, none of other 423 GV-MAGs shared >95% ANI with known reference viral genomes. When compared with previously published GV-MAGs (i.e., GOEV MAGs), additional 19 nuclecytovirus MAGs and one mirusvirus MAG produced in this study shared >95% ANI with GOEV MAGs (Table S7). It should be noted that, even when compared with the 1,065 GV-MAGs recovered from long-term sampling in Uranouchi Inlet (Fang *et al*., 2025), more than half of the 424 GV-MAGs identified in this study were missed in the seasonal sampling. Only 169 of 424 GV-MAGs exhibited ≥95% ANI with at least one of the 1,065 previously reported GV-MAGs.

**Fig. 3.**
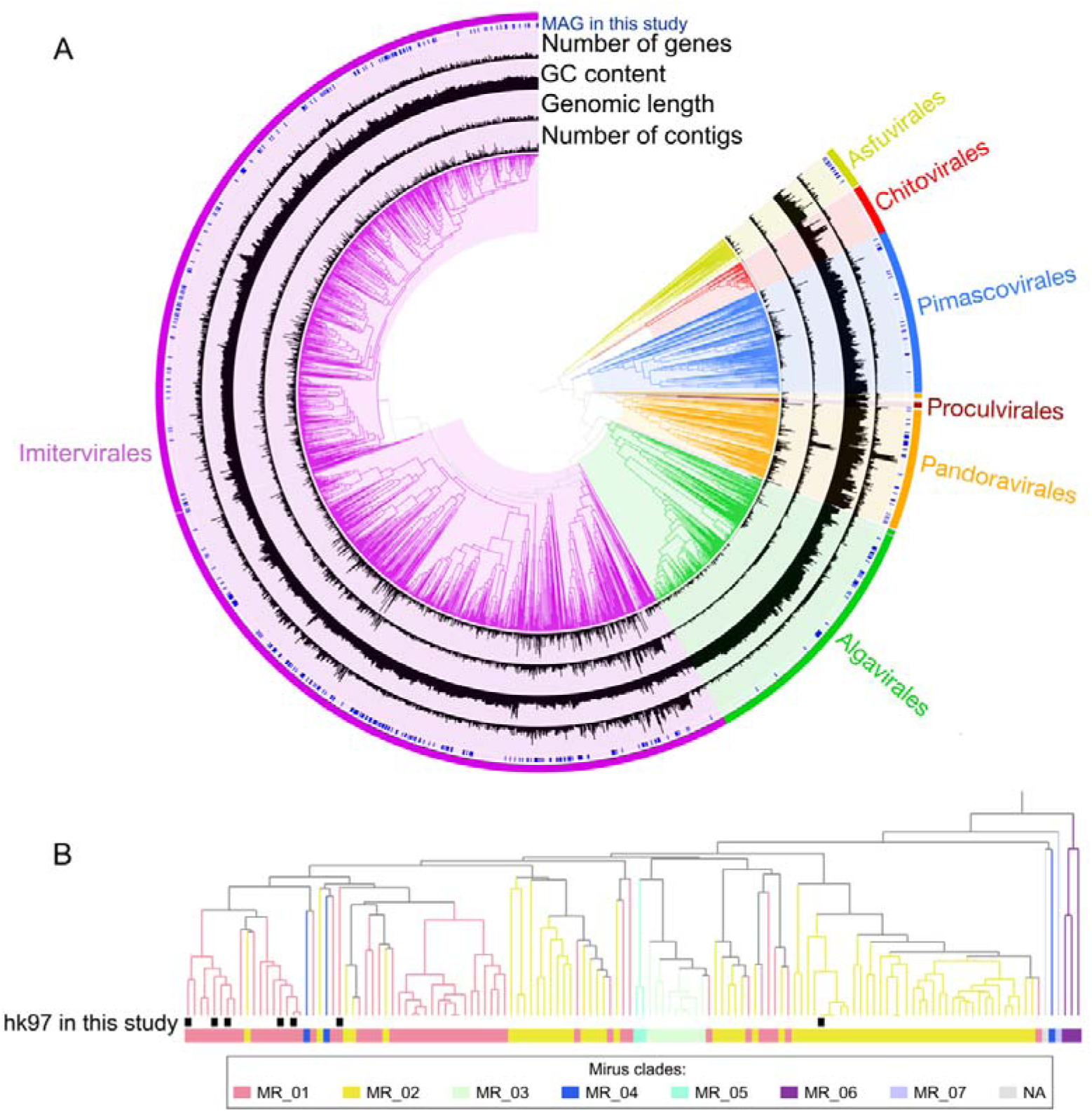
(A) Maximum-likelihood tree (1,000 bootstraps) based on 7 core marker genes of nucleocytoviruses, rooted by order *Chitovirales* and *Asfuvirales*. 1,692 GOEV nucleocytovirus genomes and 420 Uranouchi nucleocytovirus MAGs were included on the tree. (B) Maximum-likelihood tree (1,000 bootstraps) based on hk97 protein of mirusviruses, rooted by clade MR_06. 136 hk97 proteins (hmmsearch E-value<1e-5) from 77 GOEV mirusvirus MAGs and 4 Uranouchi mirusvirus MAGs were included on the tree. To facilitate unambiguous visualization of taxonomic relationships, all branches of the phylogeny were normalized to a uniform maximum evolutionary distance.

### Giant virus communities

The relative abundance of giant virus communities was dominated by *Imitervirales* (52.8% on average) in most samples, followed by *Algavirales* (28.4% on average) (Fig. S4). However, at the MAG level, HaV-MAG of *Algavirales* order was found to be most abundant in 10 samples (Fig. 1C, Table S6). In sharp contrast to the high abundance of *H*. *akashiwo* in the samples A_Day5_S and A_Day6_S, HaV-MAG remained at low relative abundance (<3%) in both samples. In the remaining 12 samples (3 A-site samples and 9 B-site samples), the relative abundance of HaV-MAG varied from 5.0% to 56.6% (24.8% on average). Besides HaV-MAG, one *Asfuvirales* MAG (Uranouchi_MAG_sc_580) showed high relative abundance (7.2% in average), whereas no other MAGs exceeded an average relative abundance of 3%. Only six GV-MAGs contributed more than 1% to the total viral community in at least one of 14 samples. Being consistent with the high relative abundance of HaV in the samples A_Day6_SCM, A_Day7_S, and B_Day3_SCM, Shannon’s diversity and evenness of the GV communities was relatively low in these samples (Fig. S3B, Table S8).

GV community structures were significantly correlated with eukaryote community structures in the B-site samples spanning 9 days (Mantel test, Pearson’s *r*=0.471, *p*<0.05), but not in the A-site samples spanning 3 days (*r*=0.076, *p*>0.05). No statistically significant separation was observed in the GV community structures between A-site (N=5) and B-site (N=9) (Fig. 2B).

### Genomic characteristics of environmental HaV

The HaV-MAG generated in this study was composed of seven contigs (258,302 bp in total), encoding 292 ORFs and displaying 30.0% GC content. Compared with the reference HaV53 genome (274,793 bp, 30.4% GC content), HaV-MAG was 16 kb shorter (Table 1). We defined ORFs shared between the two HaV genomes, when a ORF showed BLASTp best hit (E-value<1e-3) from at least one ORF from the other genome. The ORFs that did not pass this criterion was operationally denoted as ORFs specific to one of the genomes. As a result, 283 (96.9%) ORFs from HaV-MAG and 272 (93.2%) ORFs from HaV53 were found to be shared between these genomes (Table S10). A high level of collinearity and wide coverage can be also seen in Fig. S6. Some of the ORFs in HaV-MAG were incomplete when located at the edge of contigs.

**Table 1.** General features of HaV-MAG and HaV53 reference genome.

| Genome | Genome size (bp) | GC content (%) | Number of genes | Number of shared genes | Shared gene proportion | Number of specific genes | ANI |
| --- | --- | --- | --- | --- | --- | --- | --- |
| HaV-MAG | 258,302 | 30.0 | 292 | 283 | 96.9% | 9 | 99.86% |
| HaV53 | 274,793 | 30.4 | 292 | 272 | 93.2% | 20 |  |

HaV-MAG displayed the full set of core genes conserved in HaV53. Six of the nine nucleocytovirus core genes (A32, MCP, PolB, TFIIB, TopoII, VLF3) were detected in HaV-MAG, along with additional conserved genes such as TFIIS, VLTF-2, and PCNA (Table S10). Regarding predicted functions, glycosyltranferases (6 ORFs) were the most numerous (6 ORFs), followed by ring finger proteins (5 ORFs), mRNA capping enzymes (4 ORFs), KilA-N domain proteins (4 ORFs), and integrases/resolvases (4 ORFs) (Fig. S5, Table S10).

A few differences were observed in two genomes (Fig. S6). Most of the ORFs specific to HaV53 (20 ORFs) and HaV-MAG (9 ORFs) could not be functionally annotated, except transposases (3 in HaV53 and 1 in HaV-MAG), chaperone of endosialidase (HaV53), and helix-turn-helix XRE-family like protein (HaV53) (Table S9). These specific ORFs were mainly located in the 214.5 kbp–227.0 kbp region of the HaV53 genome, which is missing or largely different from HaV-MAG, as well as in the first contig of HaV-MAG, part of which is missing in HaV53 (Fig. S6). Although the chaperone of endosialidase (peptidase S74, pfam13884) in HaV53 (HaV53_86, 444 aa) was defined as specific to HaV53 by our initial analysis criterion, HaV-MAG was found to harbor a longer version of orthologous gene (HaVMAG_contig_3_1, 1529 aa) with a peptidase S74 domain (CD-search E-value: 6.36 × 10^-4^) and an additional large filamentous haemagglutinin (FhaB) domain (CD-search E-value: 1.40 × 10^-5^). FhaB is a large exoprotein domain involved in heme utilization or adhesion and was missing in HaV53_86 (Fig. S11). Another region unique to HaV53 corresponded to a repetitive coding region, predicted to encode a huge β-helical structure within an uncharacterized protein (Fig. S9) (HaV53_253: 3,663 aa; HaV-MAG_contig_6_1: 2,493 aa).

### Population structure of HaV and other giant viruses

For each GV-MAG, we calculated the number of SNV sites using the sample in which the GV-MAG displayed its maximum relative abundance. HaV-MAG exhibited a relatively large number of SNVs (3,339, in sample B_Day3_SCM). However, the nucleotide diversity of HaV-MAG was found to be relatively low (1.52×10^-3^) compared with other GV-MAGs showing a similar number of SNV sites (Fig. 4A).

**Fig. 4.**
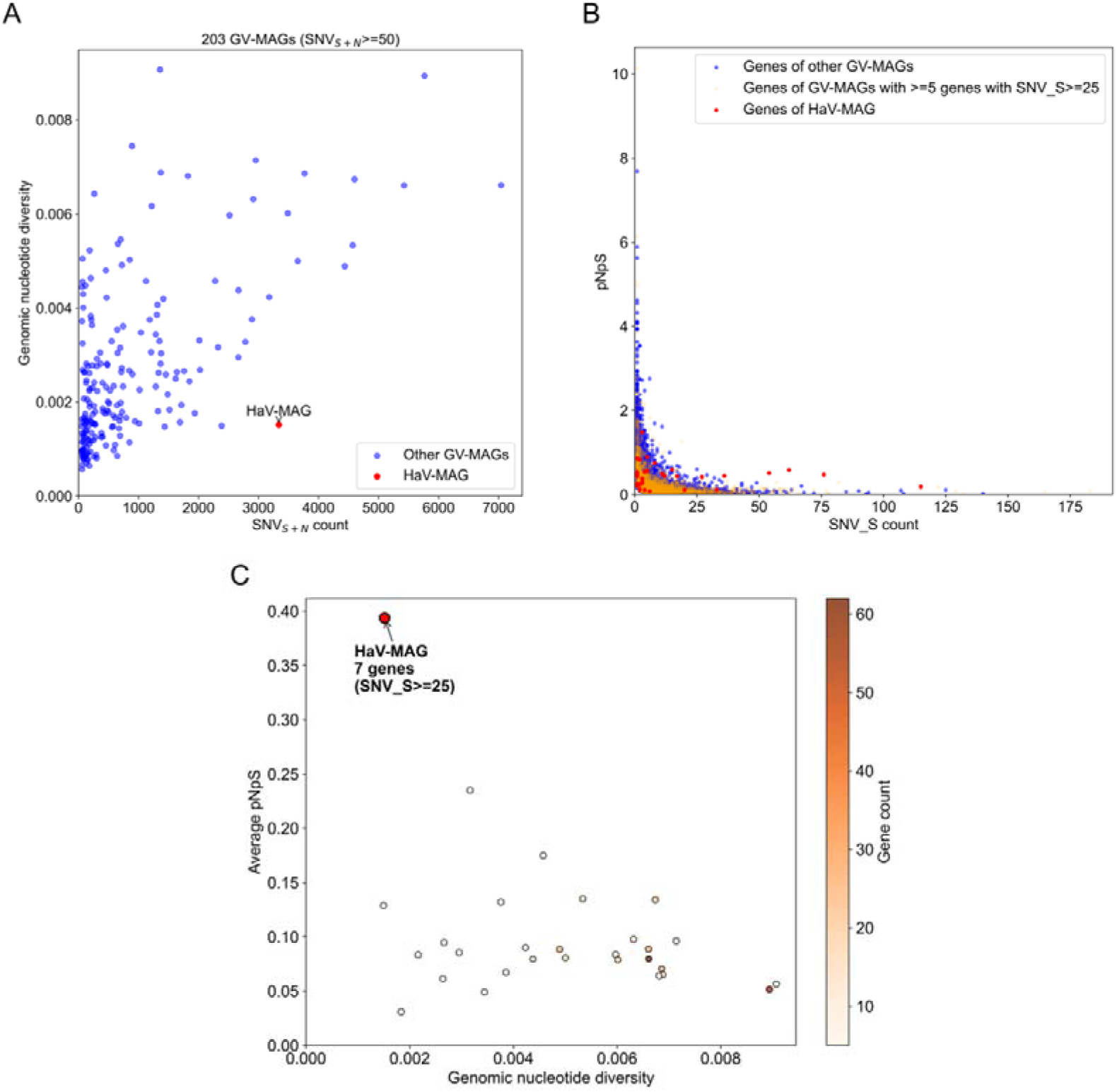
(A) Scatter plot of SNV_S+N_ count – nonsynonymous SNV count across 203 GV-MAGs with SNV_S+N_≥50. Each point represents one GV-MAG; the x-axis indicates the sum of synonymous and nonsynonymous SNVs across all genes in a MAG, and y-axis indicates its genomic nucleotide diversity. (B) Relationship between synonymous SNV count (SNV_S) and pN/pS at the gene level across 424 GV-MAGs. Genes from GV-MAGs containing ≥5 genes with SNV_S≥25 are shown in orange, genes from HaV-MAG are shown in red, and genes from other GV-MAGs are shown in blue. (C) Scatter plot of genomic nucleotide diversity versus average pN/pS across 29 GV-MAGs, each containing ≥5 genes with SNV_S≥25. Average pN/pS was calculated as the arithmetic mean of pN/pS values for genes with SNV_S≥25. All data were derived from the sample with the maximum relative abundance for each GV-MAG.

We next calculated the pN/pS ratio for each ORF in the GV-MAGs (Fig. 4B). pN/pS showed a large variance for genes with a small number of synonymous SNVs (SNV_S) (e.g., SNV_S<20), while it was more stable for ORFs with a larger number of SNV_S. This trend may be in part due to a stochastic effect for genes with a small number of SNVs. A small number of synonymous substitutions results in a highly unstable denominator, thereby leading to a large variance in the pN/pS ratio. Therefore, for small number of SNVs, the apparent elevation in pN/pS may not reflect the result of selection. Nonetheless, there was still a notable case. The ORF exhibiting the highest pN/pS ratio (1.477) in the HaV-MAG was HaVMAG_contig_3_89, which encodes a GDP-mannose 4,6 dehydratase. In this case, the ORF showed 3 synonymous SNVs and 19 nonsynonymous SNVs, which may imply a positive selection acting on this gene. As a next step, to ensure reliable estimation of pN/pS ratios, we focused on ORFs with SNV_S ≥ 25 to capture the functional constraints and purifying selection pressure in these genomes. The ORFs from HaV-MAG (7 ORFs) showed a pN/pS ratio of 0.394 on average (Fig. 4C). We then selected other 28 GV-MAGs encoding at least five ORFs with SNV_S ≥ 25 and calculated the average pN/pS ratio for each of the GV-MAGs. Of these 29 GV-MAGs, HaV-MAG displayed the highest average pN/pS ratio (Fig. 4C). Predicted functions of the seven ORFs for which pN/pS ratios were computed included a C-type lectin, a chaperone of endosialidase, a glutathione-dependent formaldehyde-activating enzyme, three helix-turn-helix domain (HTH) containing proteins and a protein of unknown function (Table S11).

### Population dynamics of HaV and other giant viruses

During the 9 days sampling at B-site, the nucleotide diversity of the HaV population represented by HaV-MAG displayed three phases with low ND compared to previous data of *Algalvirales* MAGs (Fang *et al*., 2025): (i) a gradual increase from Day-1 to Day-3; (ii) reaching a plateau from Day-3 to Day-7; (iii) a decrease from Day-7 to Day-9 (Fig. S7A). The level of the nucleotide diversity was not correlated with either the relative abundance of HaV-MAG or its host (18S relative abundance and cell density of *H*. *akashiwo*) (Pearson’s correlation test, *p*>0.05).

The genetic differences between the HaV-MAG populations in different samples were estimated using F_ST_ (253,476 comparable sites, covering 98.1% HaV-MAG). Day-2 and Day-9 populations showed relatively large genetic distances from other samples (F_ST_ on the order of 10[[), whereas pairwise F_ST_ values remained consistently low from Day-3 to Day-7 (on the order of 10[[) (Fig. S7B, C).

### Transcription landscape of HaV

At B-site, the expression levels of HaV-MAG ORFs were high within the GV communities from Day-1 to Day-8 (>3%) (Fig. 5A). Across the nine days, the highest expression levels were generally observed in a *Pandoravirales* MAG (Uranouchi_GVMAG_808). The HaV-MAG showed the second-highest average expression level overall, but on Day 3 it reached a maximum of 33% and became the most highly expressed MAG on that day. The relative transcription level of HaV-MAG was significantly correlated with the relative abundance of HaV-MAG (Pearson’s *r*=0.687, *p*<0.05) and the host *H*. *akashiwo* 18S rDNA abundance (Pearson’s *r*=0.945, *p*<0.001).

**Fig. 5.**
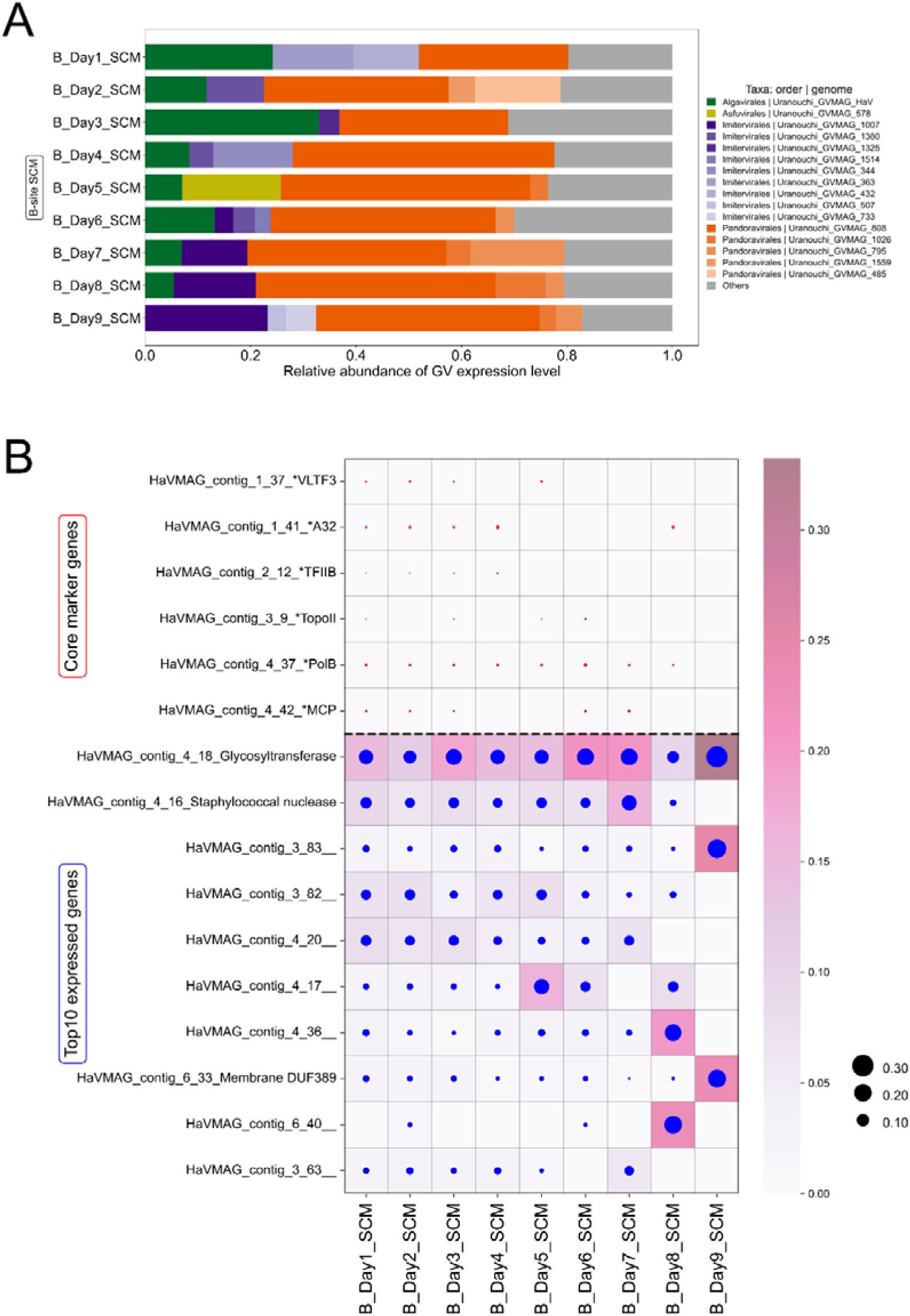
(A) Gene expression level of GV MAGs (normalized to 100% of GV community, >3% MAGs in at least one sample are listed); (B) HaV gene expression level of core marker genes and top 10 expressed genes (normalized to 100% of HaV genes).

The most highly transcribed ORF encoded a glycosyltransferase (<50% amino acid identity to any sequence in the nr database), followed by an endonuclease belongs to staphylococcal nuclease homologue (PF00565) (Fig. 5B). Although the six core marker genes of HaV-MAG were expressed from Day-1 to Day-8, their average expression levels (0.01% to 0.22% per gene) were far lower than the ten most expressed ORFs (1.46% to 15.53% per gene). The expression patterns of HaV-MAG changed with time, especially from Day-6 to Day-9 (Fig. S8).

## Discussion

Phytoplankton blooms are short-lived but high-impact events that restructure algal communities and impose strong selection on both hosts and viruses. However, how viral populations, especially those of giant viruses, evolve over the short time scales of natural blooms remains poorly resolved. *H. akashiwo* blooms often occur in late-spring and early-summer seawater (Anderson *et al*., 2002; Dursun *et al*., 2016) and have been extensively studied in the Uranouchi Inlet. In this study, our nine-day sampling campaign successfully captured a *H. akashiwo* bloom at two contrasting sites, including an innermost site (A-site) dominated by *H. akashiwo* and a site near the fishery farm (B-site) dominated by diatoms and dinoflagellates. This natural setting provided a unique opportunity to track the population dynamics of HaV *in situ*.

HaV was an abundant virus throughout this period at both A-site and B-site. Notably, HaV persisted at high relative abundance even when host densities were low (e.g., at the B-site), while conversely, low viral abundances were observed in samples with high host densities (A_Day5_S and A_Day6_S). This temporal decoupling can be explained by HaV’s long latent period (30–33 h) and large burst size, which can delay viral accumulation relative to host proliferation and sustain high viral levels after host decline (Nagasaki *et al*., 1999). A similar pattern was also reported during algal blooms in other systems (Hevroni *et al*., 2023). HaV rapidly dominated the giant virus communities following the bloom of the host population at the study site, as previously demonstrated in other coastal areas (Tomaru *et al*., 2004). The abrupt increase of HaV is thus seen as a host-specific change rather than a consequence of community-wide change.

The HaV population harbored many SNVs, but its nucleotide diversity was lower than that of other GV-MAGs with comparable numbers of SNVs (Fig. 4A). This indicates that many SNV sites in HaV-MAG were dominated by major alleles. This pattern is consistent with a genetic bottleneck or founder effect associated with the sudden initiation of infection by a limited number of viruses at the onset of host bloom, followed by explosive population expansion. During rapid expansion, new mutations might have accumulated faster than selection could efficiently filter them, potentially inflating pN/pS by allowing slightly deleterious variants to transiently persist. Furthermore, this diversification might not rely solely on *de novo* mutations; it could also reflect the rapid succession of pre-existing minor variants. Within a heterogeneous host bloom, multiple HaV genotypes that previously persisted below the detection limit may have undergone explosive proliferation upon encountering their specifically compatible host strains. Such minor variants may in part contribute to the observed microdiversity patterns. This interpretation is supported by the observation that HaV exhibited higher average pN/pS ratios than other GVs for ORFs with reliable pN/pS estimates (Fig. 4B, C). However, we cannot exclude the alternative possibility that these elevated pN/pS values result from positive selection acting on specific sequence regions of the genes used for the pN/pS calculation.

Temporal dynamics of HaV population structure further support the view of genetic bottleneck. The nucleotide diversity of the HaV population increased during the early phase of the sampling period, which could reflect either the ongoing generation of genetic diversity during viral proliferation, or the simultaneous expansion of pre-existing minor viral genotypes from environmental reservoirs (such as sediment) or background populations that were previously rare. Subsequently, diversity decreased toward the end of the sampling period (Fig. S7A). This decline in viral microdiversity suggests a limited number of successful genotypes in infecting available host. However, because it is difficult to ensure that identical water masses were sampled throughout the time series, these fine-scale dynamics should be interpreted with caution.

At the functional level, several results indicated that the host-virus interaction is the critical factor shaping the HaV population during rapid population expansion. Transcriptomic analysis showed that a HaV-specific glycosyltransferase was expressed at much higher levels than core genes (Fig. 5B), in contrast to previous observations in which GV transcription was strongly biased toward core genes (Ha *et al*., 2021). GTs encoded by giant viruses are known to modify viral or host glycoproteins by manipulating sugar residues, thereby facilitating host recognition, attachment, and immune evasion (Speciale *et al*., 2022). The HaV GT belongs to the non-Leloir GT-C fold (Mestrom *et al*., 2019) and contains multiple predicted transmembrane α-helices (Fig. S10), indicating that it is likely involved in glycosylation of viral capsid or surface-associated proteins. Therefore, the high expression of GT may enhance HaV infection efficiency through glycosylation-mediated adsorption or by modifying viral components to evade host antiviral mechanisms. Consistent with this interpretation, infection specificity in the HaV–*H. akashiwo* system has been linked to strain-specific differences in viral adsorption to host cell surfaces (Funaoka *et al*., 2023).

An additional perspective on rapid diversification at the host-virus interface is provided by gene gain and loss (Table S9). Transposases, which serve as the key enzymes mediating DNA excision and transfer (Sun *et al*., 2015), were found to be absent from the HaV-MAG but present in the reference HaV53 genome. Although we could not confirm whether the divergent transposases originate from a single transposable element, their heterogeneity suggests variable patterns of DNA fragment transfer during interactions with diverse *H*. *akashiwo* strains. In addition, the viral chaperone of endosialidase, which is essential for the assembly and folding of endosialidases that degrade surface sialic acids used by hosts for pathogen recognition (Schwarzer *et al*., 2007; Tiralongo, 2010). Specifically, the orthologous gene in HaV-MAG (HaVMAG_contig_3_1, 1529 aa) contains an additional FhaB domain that is absent in HaV53_86 (444 aa), suggesting a possible “domain gain” or “loss” event in HaV genomes during population evolution (Table S9; Table S10; Fig. S11). Divergence in these surface- and interaction-related genes may modulate infectivity toward hosts exhibiting distinct extracellular defense traits. Together, these observations suggest that gene gain and loss contribute to maintaining a selective advantage in the HaV-host arms race.

For the first time, our study resolved the genetic structure, population dynamics, and transcriptional activity of a natural HaV population. The genetic diversity of the HaV population was found to be low, but it transiently increased during the proliferation period associated with the host bloom. This dynamic genetic shift suggests that the HaV population rapidly adapts to host blooms through a combination of *in situ* evolution and the selection of standing genetic variation. In parallel, environmental HaV genomes differ in gene content from the reference HaV53 genome, indicating genomic plasticity of HaV in natural populations. Functionally, HaV exhibits low expression of core genes but high expression of accessory genes, involved in host-specific interactions or evasion of host defenses. Together, these observations support host–virus interactions acting as purifying selective pressure on HaV during the bloom.

## Acknowledgements

This study was supported by JSPS/KAKENHI (nos. 18H02279 and 19H05667 to Hiroyuki Ogata, 17H03850 and 21H05057 to Takashi Yoshida and Hiroyuki Ogata, and nos. 19K15895 and 23K23685 to Hisashi Endo), Scientific Research on Innovative Areas from the Ministry of Education, Culture, Science, Sports and Technology (MEXT) of Japan (nos. 16H06429, 16K21723, and 16H06437 to Hiroyuki Ogata), JST PRESTO (no. JPMJPR23G3 to Hisashi Endo), and the Collaborative Research Program of the Institute for Chemical Research, Kyoto University (grant numbers 2016-28 and 2019-35). We would like to express sincere gratitude to the staff of SuperComputer System, Institute for Chemical Research, Kyoto University. We thank Kochi Prefecture Fisheries Promotion Department for the sampling permission and providing information of algal bloom area, weather and hydrological characteristics. We also would like to thank Florian Prodinger, Chi-Yu Shih, Wenwen Liu, Tatsuhiro Isozaki, Hiroaki Takebe, Kento Tominaga and Yoshiaki Sato for the kind instructions in both dry-lab and wet-lab experiments.

## Supplementary figures and text

**Fig. S1.**
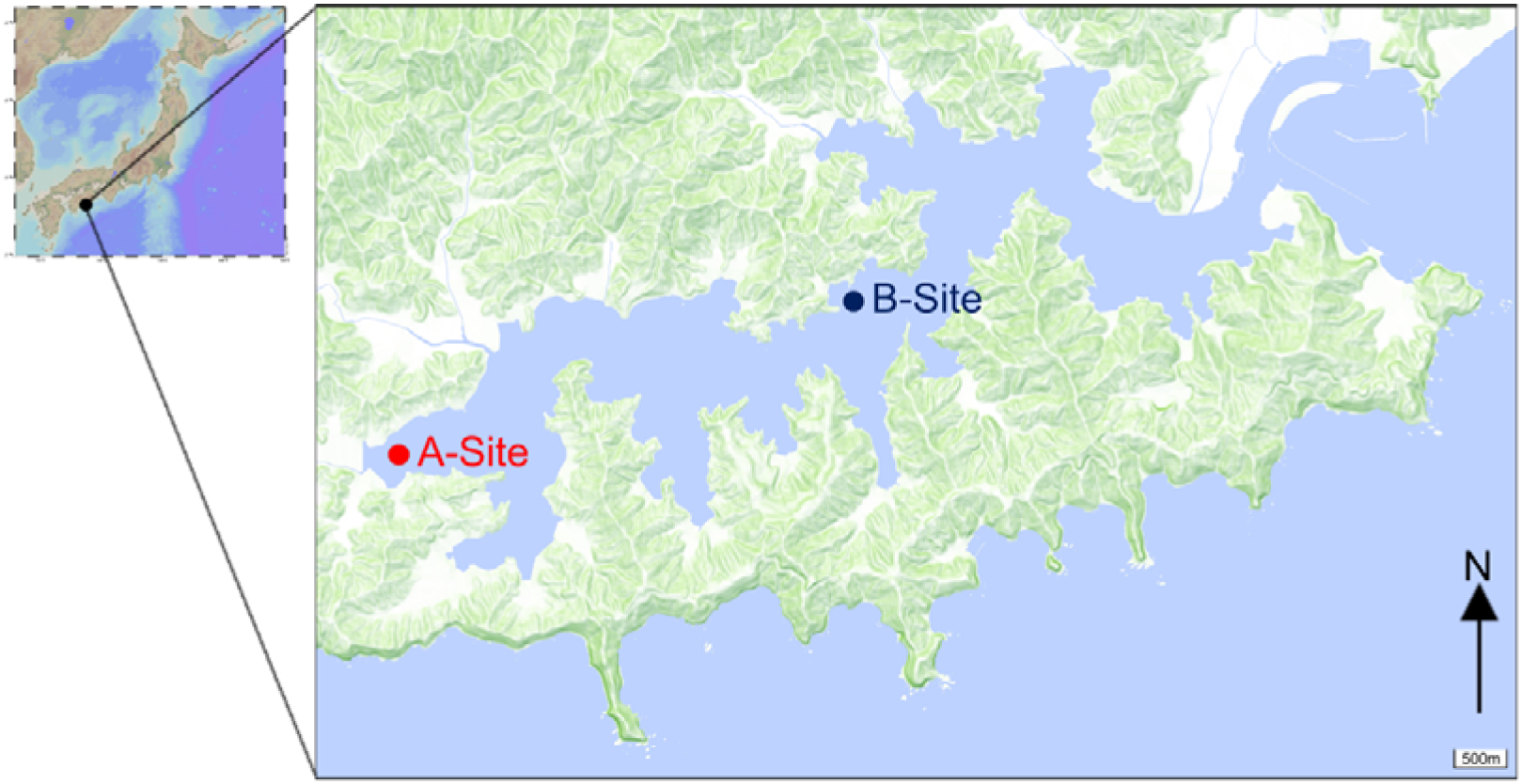
Location of Uranouchi Bay and the map of sampling sites in this study (A-Site in red tide area of innermost bay [IMB] with red mark, and B-Site closed to a fishery farm of western middle bay [WMB] with deep-blue mark). Base map data are provided by the Ocean Data View (ODV, version 5.7.2) and Geospatial Information Authority of Japan (GSI Maps Vector).

**Fig. S2.**
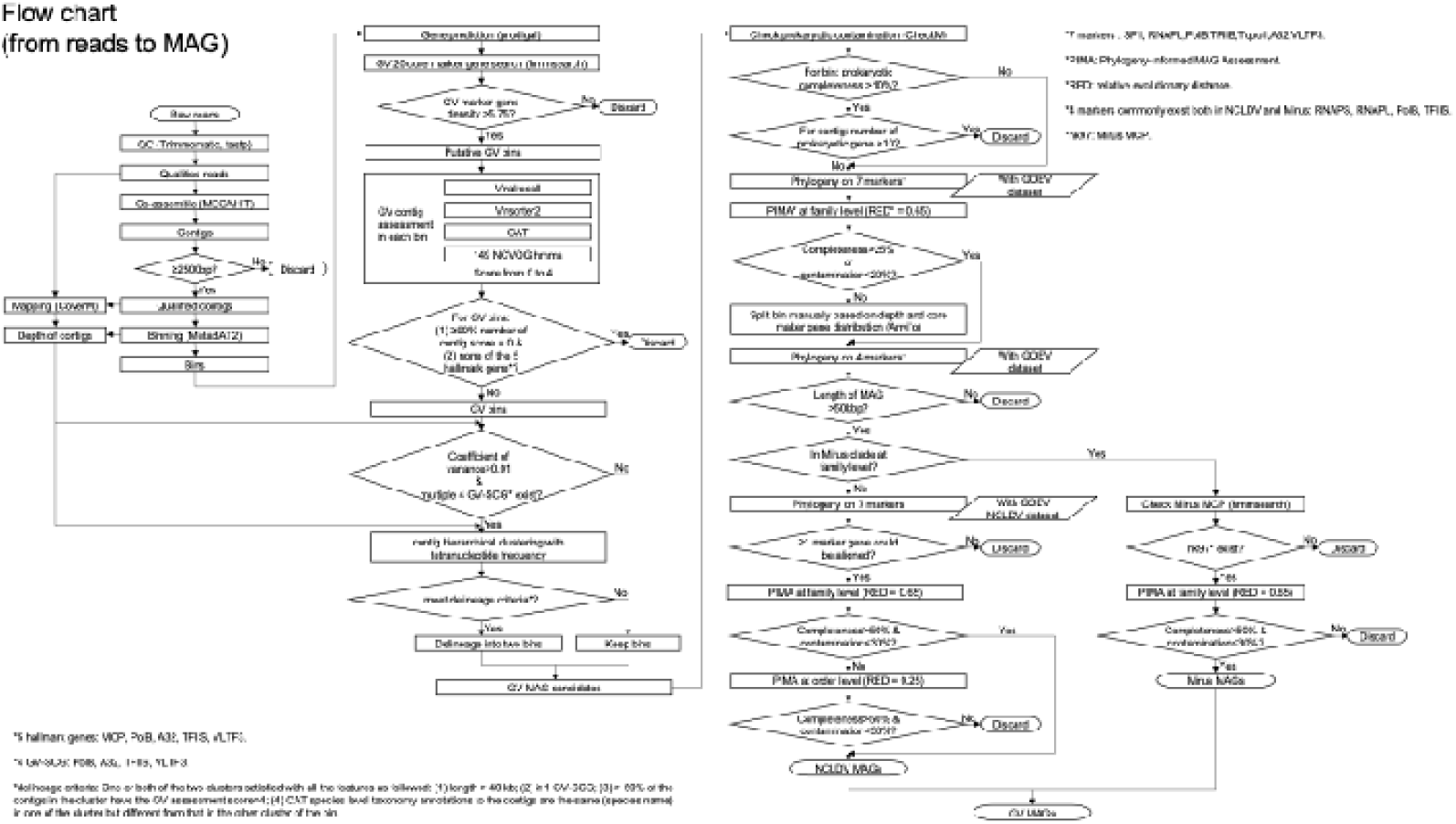
Flow chart of GV-MAG generating pipeline. (see **Fig. S2 supplementary text** below)

**Fig. S3.**
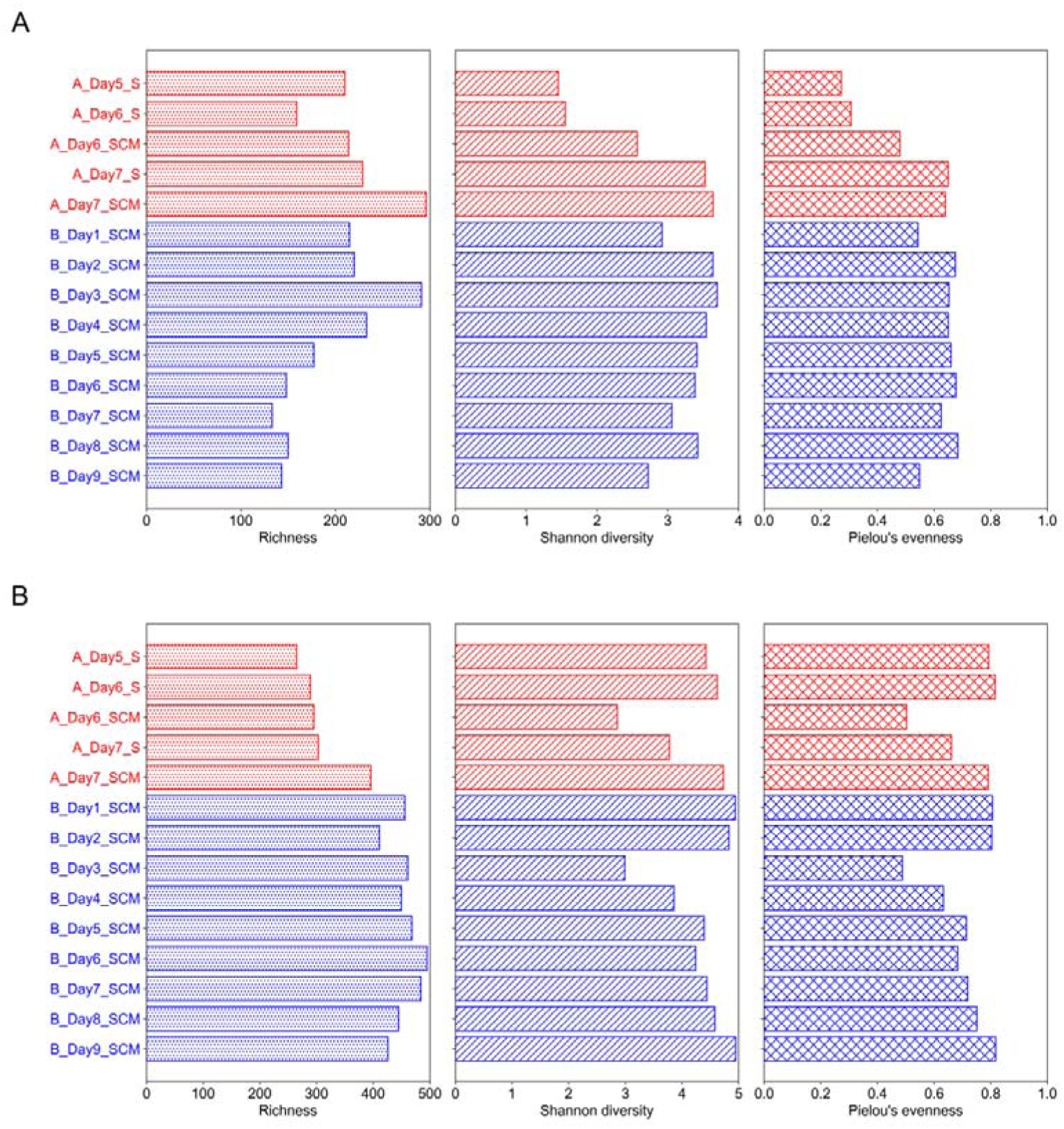
Richness, Shannon diversity and Pielou’s evenness of 14 samples in this study: (A) 3–144 µm microbial eukaryotic 18S community (non-singleton ASVs without metazoa or fungi); (B) 0.22–3 µm giant virus community. S: surface; SCM: Subsurface Chlorophyll Maximum.

**Fig. S4.**
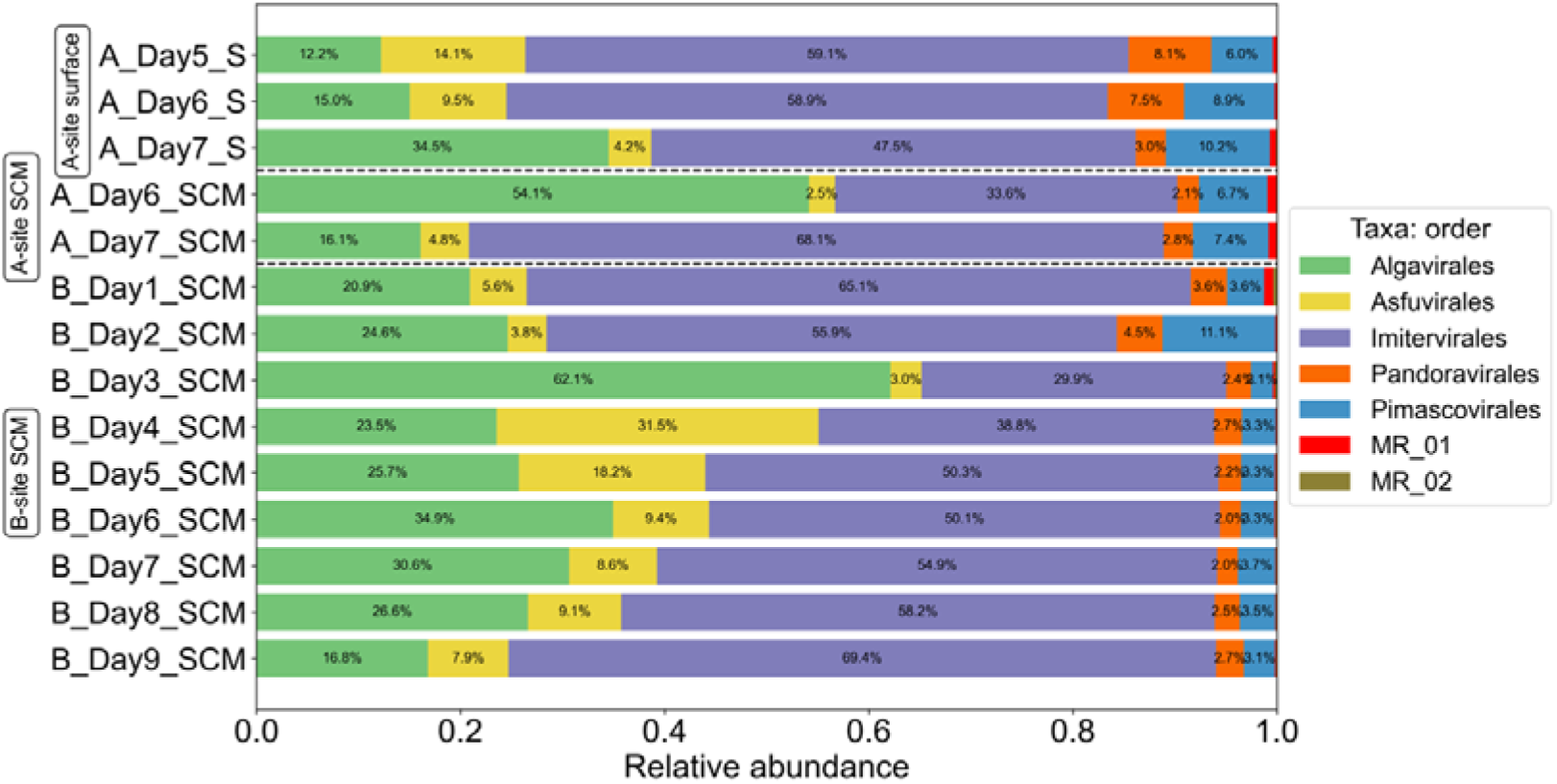
Relative abundance of GV-MAGs and GOEV genomes in order level in the 14 samples.

**Fig. S5.**
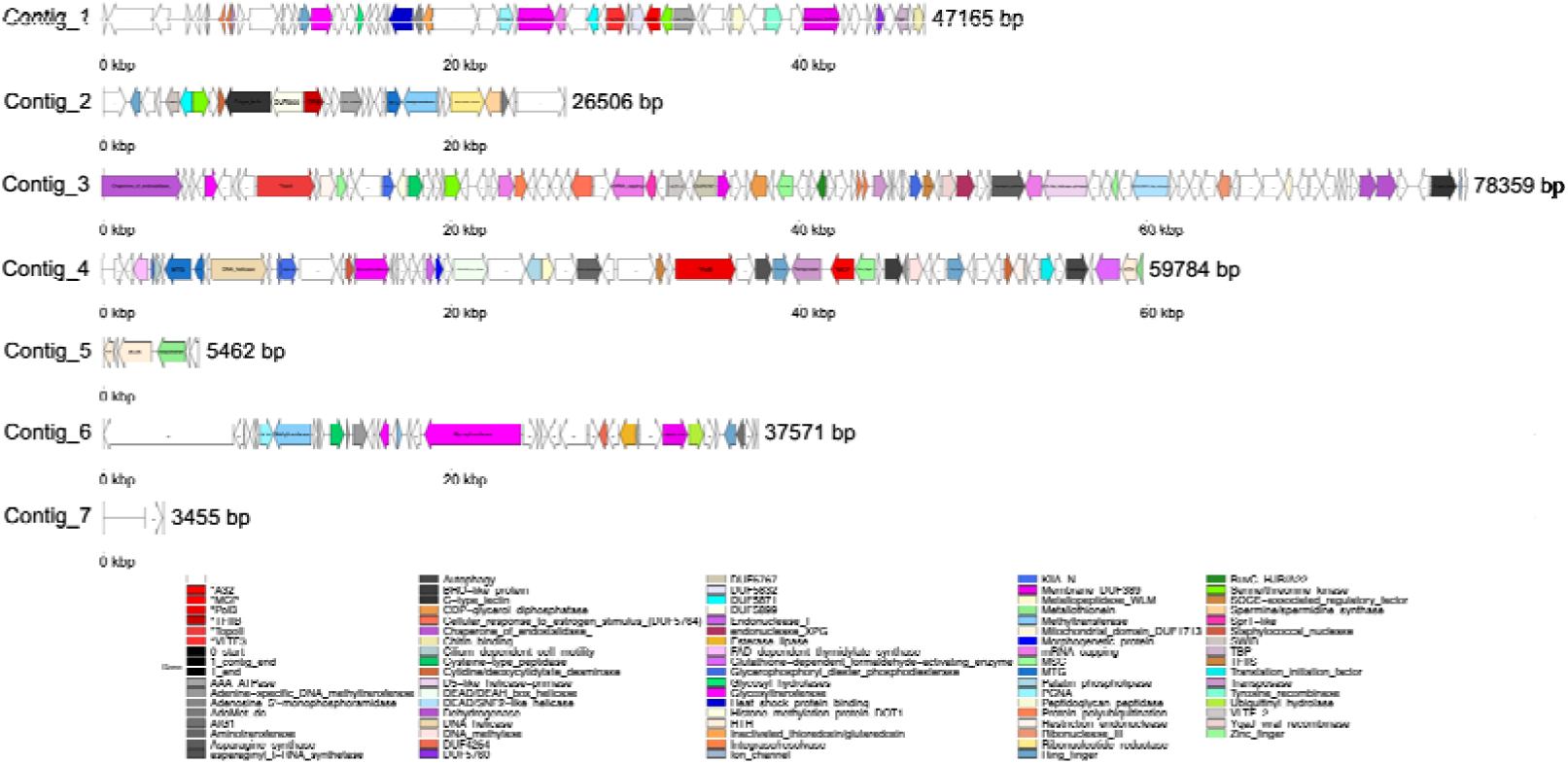
Genomic structure of HaV-MAG in this study. “*” marked gene refers to one of the nine core marker genes of nucleocytoviruses.

**Fig. S6.**
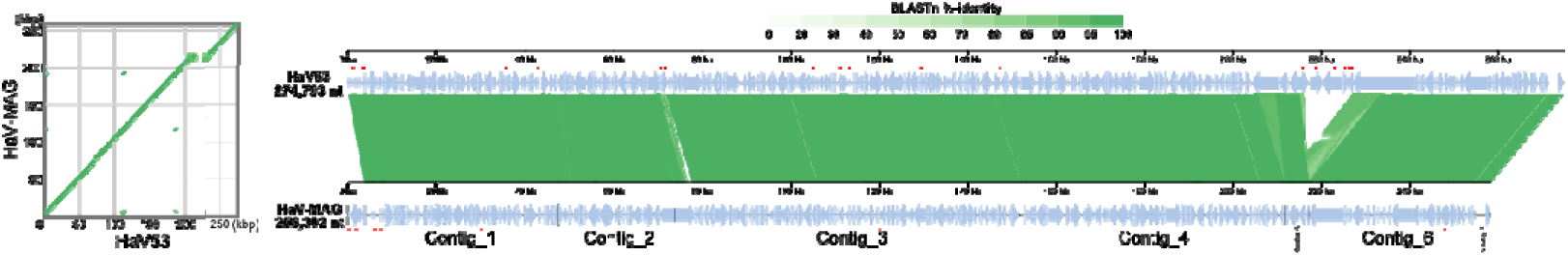
Left: synteny map of HaV53 and HaV-MAG; right: pairwise gene alignment between HaV53 and HaV-MAG. The nucleotide dissimilar ORFs were marked with red small dot on the right figure (linked to Table S9).

**Fig. S7.**
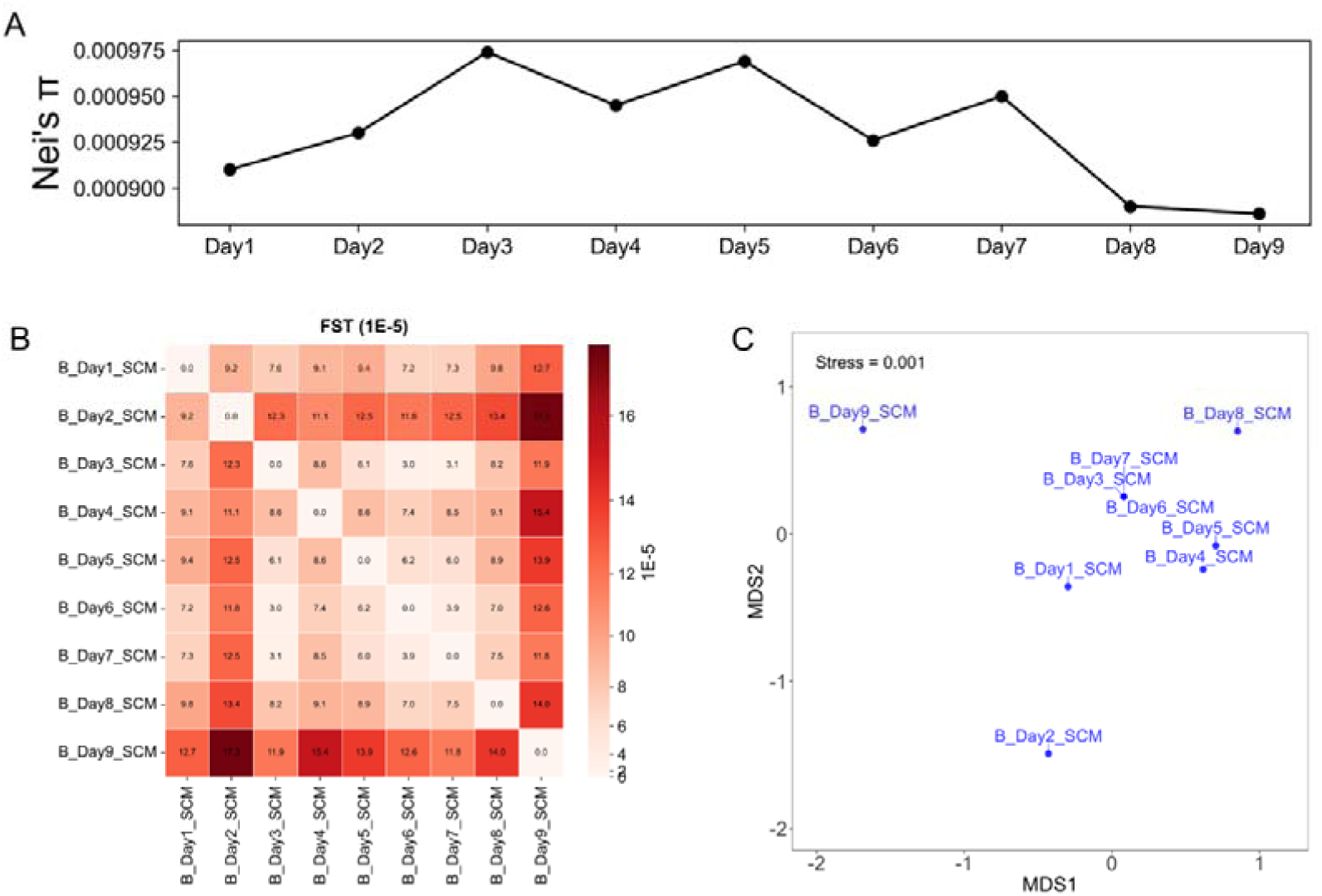
(A) Nei’s nucleotide diversity (Nei’s π) of HaV in B-Site during the 9 days. (B) Fixation index (F_ST_) heatmap of HaV population in B-Site SCM samples, which is from 0 (totally the same) to 1 (totally different). The numerical values displayed in each cell equal the original F_ST_ estimates multiplied by 10^5^. (C) NMDS of F_ST_ among HaV 9-days population in B-Site.

**Fig. S8.**
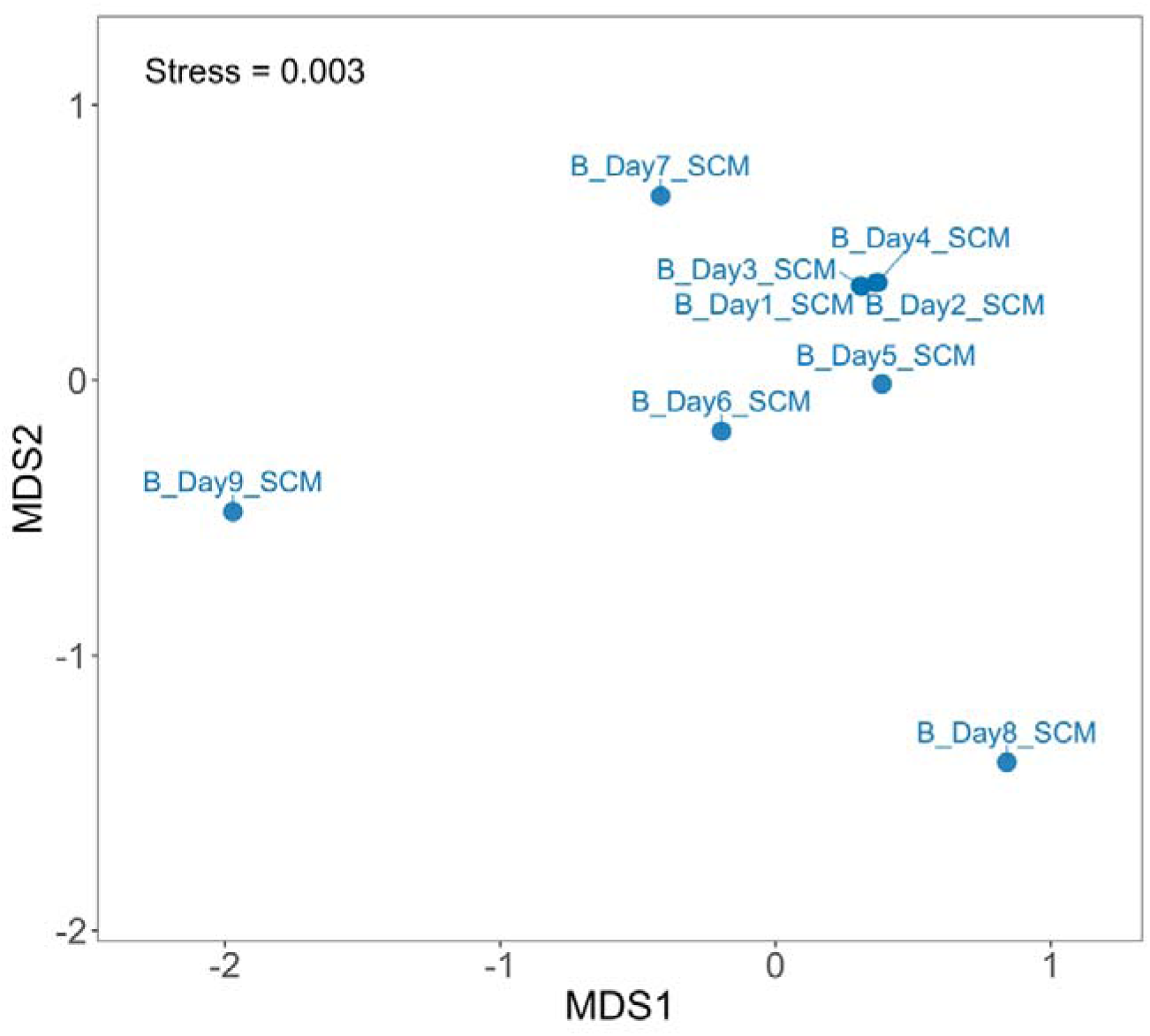
Dynamics of relative abundance of HaV gene expression level in B-Site 9-day samples.

**Fig. S9.**
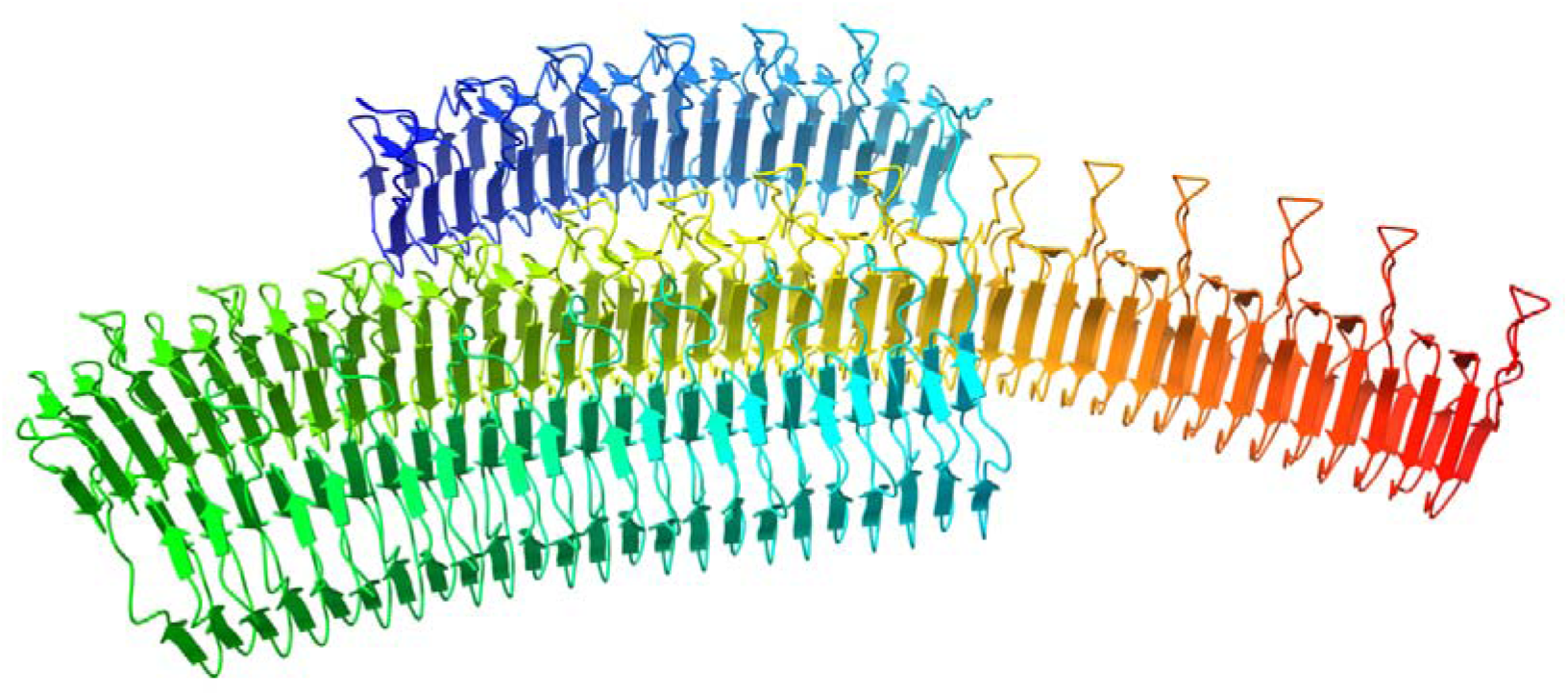
Partial (from 2580 aa to 4663 aa) structure of the huge protein HaV53_253, the missing repeat-rich region in protein HaVMAG_contig_6_1. N-terminal to C-terminal were colored from red to blue.

**Fig. S10.**
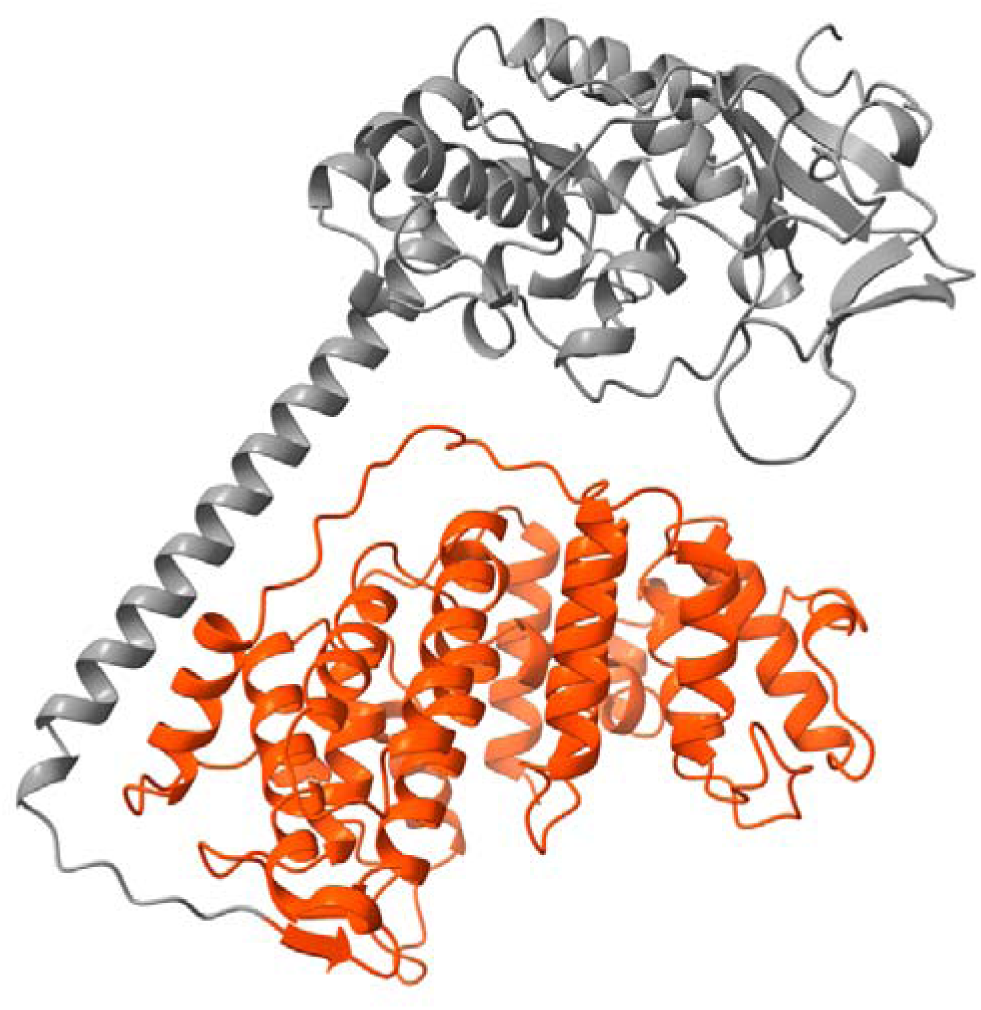
Protein structure of highly expressed HaV specific glycosyltransferase (GT) in HaV-MAG (HaVMAG_contig_4_18). The α-helices rich region which could define this GT-C fold is highlighted with orange color.

**Fig. S11.**
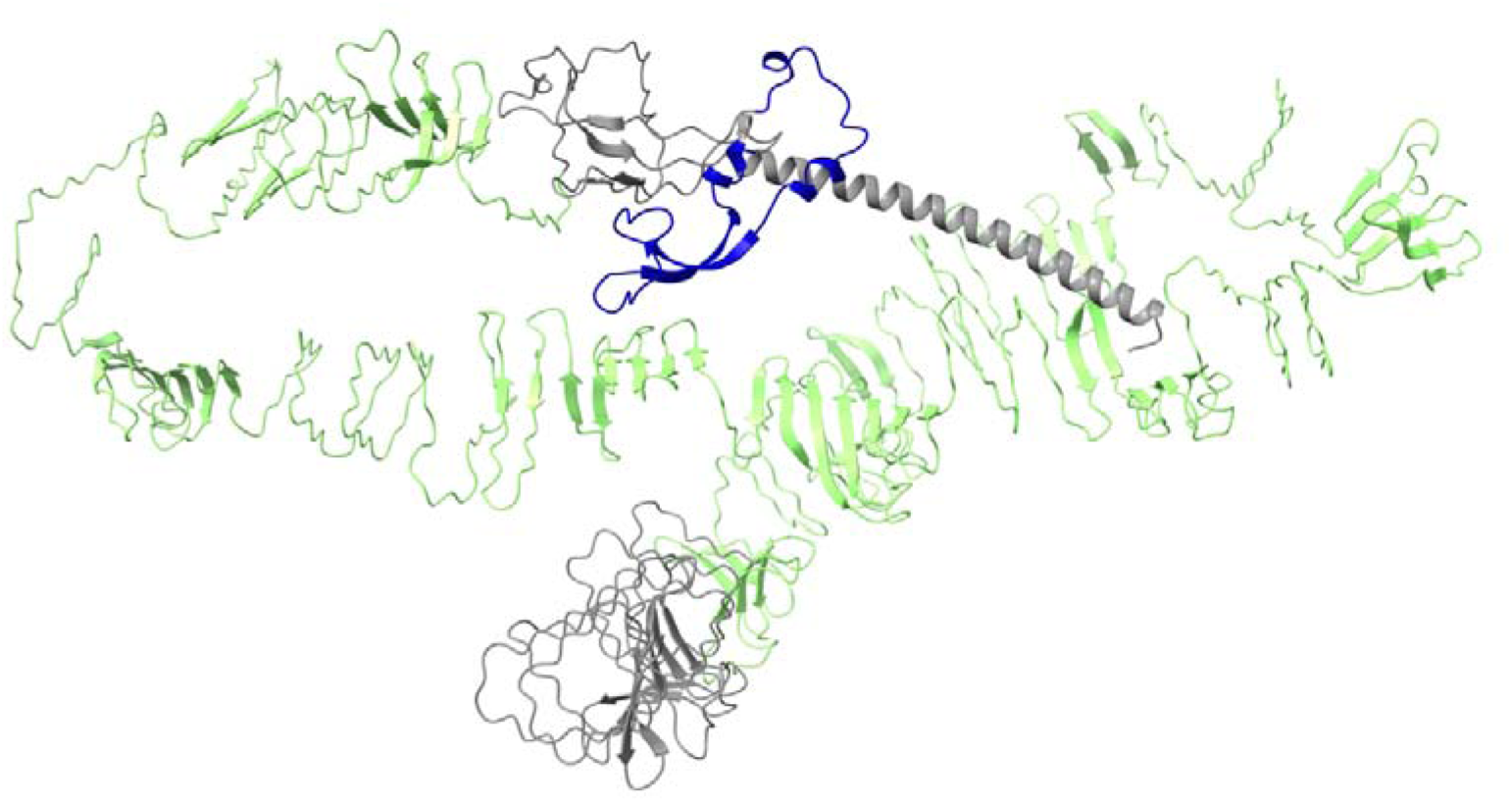
Protein structure of HaVMAG_contig_3_1 (1530aa) in HaV-MAG. It contains a specific chaperone of endosialidase (deep blue, residues 1380–1439) and a large FhaB domain (light green, residues 217–1329).

## supplementary text

PART I: reads QC, assemble, initial binning. Reads quality control was performed by Trimmomatic and fastp, then co-assembled by MEGAHIT for samples in both size-fractions. Contigs with length more than 2500 bp were kept for downstream analysis. Depth of each contigs were calculated based on qualified reads mapping on contigs by CoverM (v0.7.0) (Aroney *et al*., 2025). Binning was performed by MetaBAT2 (v2.15) (Kang *et al*., 2019).

PART II: selection of GV-MAG candidates. Predicted ORFs (by prodigal [v2.6.3] [Hyatt *et al*., 2010]) in total bins were matched on 20 core marker gene Hidden Markov models (hmms) of giant viruses by hmmsearch. Bins with GV core gene density over 5.75 were kept as putative GV bins (Fang *et al*., 2025). GV contig assessment was performed sequentially by ViralRecall (v2.1) (Aylward and Moniruzzaman, 2021), VirSorter2 (v2.2.3) (Guo *et al*., 2021), CAT database (v2021-01-07) (von Meijenfeldt *et al*., 2019), 149 NCVOG hmms (Yutin *et al*., 2009; Gaïa *et al*., 2023). Each passed step will make the bin score “+1”. A bin was treated as unqualified when both two conditions met: (1) over 90% number of contigs scored as 0; (2) none of the five hallmark genes (MCP, PolB, TFIIB, TopoII, and A32) found. While depth coefficient of variation >0.01 and multiple copies of 4 GV-SCGs exist, a bin was further delineaged into two bins based on tetranucleotide frequency and depth.

PART III: GV-MAG refinement. Prokaryotic completeness in each bin was calculated by CheckM (v1.2.2) (Parks *et al*., 2015). If contamination >10% and number of prokaryotic gene >10 in a bin or concentrated in a contig, it was discarded. Nucleocytovirus or mirusvirus core marker gene sets in protein format were searched, concatenate-aligned and trimmed by ncldv_markersearch.py (v1.1) (Aylward *et al*., 2021), followed with phylogenetic analysis by iqtree2 (v2.2.2.6) (Minh *et al*., 2020). Giant virus genome completeness and contamination were evaluated by Phylogeny-informed MAG assessment (PIMA) (Fang *et al*., 2025). The bins with completeness >25%, contamination <20% and size >50 kbp were then kept. MAG refinement based on contig depth Dendrogram was manually performed by Anvi’o (v7.1) (Eren *et al*., 2015). Final nucleocytovirus MAGs should be qualified if (1) on the 7 core marker gene tree; (2) completeness >50% and contamination <30% at both family level (RED>0.65) and order level (0.25<RED≤0.65). For recognizing Mirus features, conditions below should be satisfied in a MAG: (1) contained in Mirus families; (2) Mirus-specific major capsid protein hk97 (Gaïa *et al*., 2023) detected (hmmsearch [v3.4] [Finn *et al*., 2011], E-value<1e-5); (3) completeness >50% and contamination <30%.

## Notes

https://www.genome.jp/ftp/db/community/Uranouchi_GVMAGs_9days/

### Competing Interest Statement

The authors have declared no competing interest.

